# Neuronavigation-free and MRI-free Localization of Deep Brain Targets

**DOI:** 10.64898/2026.02.04.703768

**Authors:** Yishai Valter, Yu Huang, Niranjan Khadka, Abhishek Datta, Marom Bikson

## Abstract

Emerging non-invasive brain stimulation modalities, including transcranial focused ultrasound and transcranial interferential stimulation, offer the promise of safely and non-invasively modulating deep brain structures. Accurate targeting of these regions typically involves subject-specific MRI, limiting widespread deployment. Here, we introduce an MRI-free, neuronavigation-free method for localizing deep brain targets using only three simple scalp measurements. These measures define an affine transformation that maps the MNI152 standard head model to an individual’s head geometry. We evaluated our approach on 50 healthy adults, comparing our model◻predicted coordinates against ground-truth coordinates obtained via MRI-based nonlinear normalization. Across ten deep brain targets, our method achieved a mean localization error of 3.82 mm demonstrating a more accessible and cost-efficient alternative than MRI- or neuronavigation-based approaches.

## Introduction

Emerging noninvasive brain stimulation modalities such as transcranial focused ultrasound (tFUS) and transcranial temporal interference (TI) stimulation can noninvasively modulate deep brain structures. Accurate localization of these techniques, however, typically depends on subject-specific magnetic resonance imaging (MRI) (Truong et al., 2022; Brahma et al., 2025).

Riis et al. (2025) introduced an MRI-free, neuronavigation-based method for deep brain localization by fitting the MNI152 template to subject-specific head geometry based on facial landmarks and found a mean localization error of 5.75 mm. Nafziger et al. (2025) further reduced the mean localization error to 3.9 mm by co-registering 25 manually identified scalp landmarks to the MNI152 template. These approaches depend on neuronavigation systems with associated cost, complexity, and portability limitations and depend on operator time for manual localization of numerous scalp points with associated errors.

Several MRI-free, neuronavigation-free methods have been developed to localize scalp targets, such as Beam-F3 (Beam et al., 2009), EPlacement (Fabregat-Sanjuan et al., 2023), continuous proportionate coordinates (Jiang et al., 2022), and tetra codes (Li et al., 2024). Beam-F3 has especially proven useful in transcranial magnetic stimulation for locating the scalp region above the left dorsolateral prefrontal cortex (Valter et al., 2024) using only three gross anatomical measurements and a democratized algorithm. However, because these methods rely on scalp coordinate mapping rather than volumetric modeling, their applicability is inherently limited to superficial cortical regions.

We recently introduced a volumetric modeling approach that estimates individualized head geometry using only three anatomical measurements adapted from the Beam-F3 method: head circumference, nasion-to-inion distance, and tragus-to-tragus distance. These measurements are used to derive a scaling matrix which is then applied to the MNI152 template, yielding a subject-specific volumetric head model. The method was validated *in silico* for cortical target localization (Valter et al., 2025) and made available in a web-accessible format (tmstargets.com).

In the present study, we evaluate the utility of the Valter 2025 approach for deep brain target localization. Using MRI data from fifty participants, we compared the coordinates of ten deep brain targets predicted by our scalp measurement-based approach with those obtained via MRI-based nonlinear normalization and found a mean localization error of 3.82 mm, representing comparable accuracy to a previously reported neuronavigation-based method (Riis et al., 2025). To contextualize these results, we further compared localization errors from MRI-based linear normalization and from the unscaled MNI152 template. MRI-based nonlinear localization served as the ground truth for all comparisons.

## Methods

### MRI Dataset

We retrieved the first fifty MR scans from a publicly available dataset consisting of healthy adult volunteers (Reddan et al., 2025) hosted on OpenNeuro.org. Three scans were excluded due to partial omission of the ears, which prevented measurement of the tragus-to-tragus distance. These scans were replaced with the next available subjects in the dataset to maintain a total of fifty data sets.

### Brain Targets

Ten deep brain targets were selected for analysis. Seven of these corresponded to those reported by Riis et al. (2025) for neuronavigation-based targeting including three sites within the subgenual anterior cingulate cortex (SGC; Brodmann area 25) and four sites spanning the pregenual to anterior midcingulate cortex (PGC, aMCC; Brodmann areas s24, p24, a24, and 33). These targets lie within the midsagittal plane (y–z axes) at approximately 5 mm intervals, facilitating systematic evaluation of targeting accuracy along a deep midline trajectory. An additional three targets—right hippocampus [28, −22, −14], right amygdala [21, −1, −22], and right caudate [14, 13, 11]—were included from Brahma et al. (2025) due to their relevance as common deep brain stimulation sites. In total, ten MNI-defined targets were analyzed (Table 1).

**Table 1.**
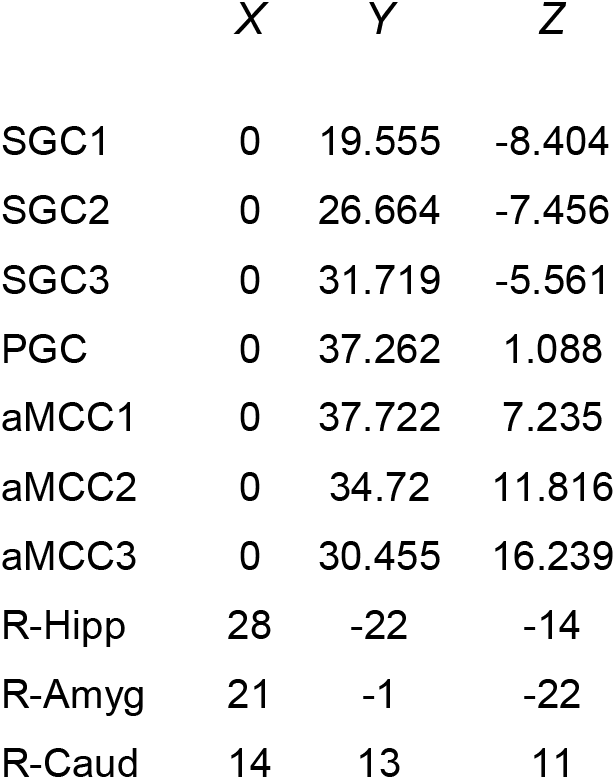
Ten deep brain targets were selected for analyzing localization accuracy. Each target was identified by its MNI coordinates. (SGC: subgenual anterior cingulate cortex, PGC: pregenual midcingulate cortex, aMCC: anterior midcingulate cortex)

|  | X | Y | Z |
| --- | --- | --- | --- |
| SGC1 | 0 | 19.555 | -8.404 |
| SGC2 | 0 | 26.664 | -7.456 |
| SGC3 | 0 | 31.719 | -5.561 |
| PGC | 0 | 37.262 | 1.088 |
| aMCC1 | 0 | 37.722 | 7.235 |
| aMCC2 | 0 | 34.72 | 11.816 |
| aMCC3 | 0 | 30.455 | 16.239 |
| R-Hipp | 28 | -22 | -14 |
| R-Amyg | 21 | -1 | -22 |
| R-Caud | 14 | 13 | 11 |

Brain extraction was performed on the subjects’ scans using FSL’s brain extraction tool (*bet*). Extraction failed for one subject, whose data were replaced with the next available scan. Four different localization methods were then performed: 1) MRI-based linear localization, 2) MRI-free, neuronavigation-free, scalp measurement-based localization, 3) unscaled MNI template, and 4) MRI-based nonlinear localization (Figure 1).

**Figure 1.**
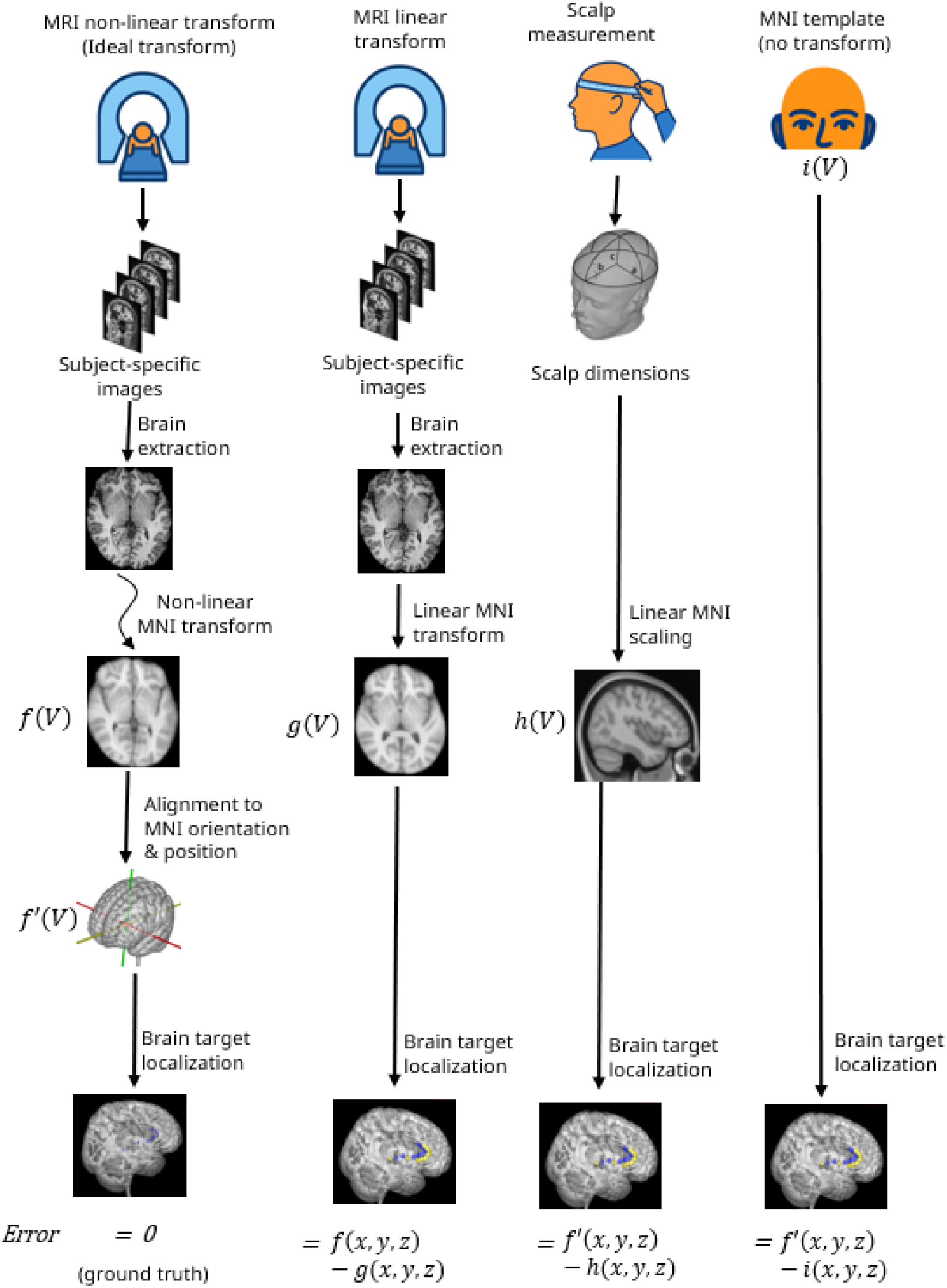
Four localization methods were performed: MRI-based nonlinear normalization (ground truth), MRI-based linear normalization, MRI-free, neuronavigation-free scaling using scalp measurements, and the unscaled MNI template. Error was quantified as the Euclidean distance between the predicted coordinates and the coordinates defined by the MRI-based nonlinear localization (ideal transform / ground truth). *f* denotes the nonlinear transformation and *f’* denotes the nonlinear transformation after rigid body alignment to MNI orientation and position. MRI-free localization accuracies were tested using *f’* as the ground truth.

### 1. MRI-based linear localization

For each subject, an affine transform was applied to the MNI brain using the subject’s brain as the reference image (FSL, *flirt*). The transforms were saved and applied to the target coordinates in template space (FSL, *img2imgcoords*) to obtain subject-specific coordinates for each brain target.

### 2. MRI-free, neuronavigation-free, scalp measurement-based transform

The subject’s scans were converted from NIfTI to DICOM format using 3D Slicer (slicer.org) and imported into Horos (horosproject.org). Three virtual scalp measurements were obtained using the curvilinear measurement tool: head circumference, nasion-to-inion distance, and tragus-to-tragus distance. These measurements were used to approximate a scaling matrix according to the method previously described (Valter et al., 2025). The derived scaling matrices were then applied to the target coordinates in template space to obtain subject-specific coordinates for each brain target.

### 3. MNI template unscaled

We also considered the unscaled MNI template to explore overall differences in deep brain targets between each individual brain and a standard template. For this method, subject-space is assumed to be the same as template space, so any brain target is predicted to be at its MNI coordinates.

### 4. MRI-based nonlinear localization

Nonlinear transformations were generated (FSL, *fnirt*) using each subject’s volume as the reference image and their respective linear transform matrix as the initial estimate. This nonlinear registration served as the ground truth as it provides the most anatomically accurate MNI-to-subject mapping by warping the MNI volume to individual anatomy. The resulting warp fields were applied to the target coordinates in template space (FSL, *img2imgcoords)* to obtain subject-specific coordinates for each brain target.

To perform a meaningful comparison between the MRI-free methods (methods 2 and 3) with the ground-truth nonlinear MRI localization (method 4), all results needed to be placed in the same orientation. We did this by identifying three midsagittal points in each subject’s brain using the nonlinear warp field and the *img2imgcoords* tool: the anterior commissure (AC; [0, 0, 0]), the posterior commissure (PC; [0, −25, 0]), and a third point on the midline ([0, 0, 50]). Using these three points, we computed a rigid-body transformation that aligned each subject’s AC-PC line with the y-axis and fixed the y-axis rotation using the third point. Because rigid-body transforms adjust only position and orientation, this step preserved each subject’s anatomy while placing all brains into a common coordinate frame. We then applied this transform to the nonlinear-derived coordinates, producing aligned subject-specific coordinates that allowed direct comparison with the MRI-free predictions.

### Error Analysis

Localization error was quantified as the Euclidean distance between each target coordinate derived from the experimental methods (1, 2, and 3) and the MRI-based nonlinear transformation (method 4; ground truth). To assess relative performance, paired t-tests were conducted to compare localization errors across the three experimental approaches.

## Results

Across the ten targets, mean errors for MRI-based linear localization ranged from 2.66 mm to 4.57 mm with a mean of 3.59 ± 1.78 across all targets (Figure 2). This method provided significantly greater accuracy than other methods for targets within the aMCC (p-values ranging from 0.02 to 0.05), but not for other targets. Mean errors for scalp-based MRI-free localization ranged from 2.66 mm to 4.57 for each of the ten targets with a mean of 3.82 ± 1.78 across all targets. Accuracy for scalp measurement-based localization was found to be significantly greater than localization using the unscaled MNI template for targets within the aMCC, right caudate, and right hippocampus (p-values ranging from <0.001 to 0.02). Mean errors for the unscaled MNI method ranged from 2.93 mm to 5.98 mm for each of the ten targets with a mean of 4.55 ± 1.74 across all targets. Surprisingly, this method provided the greatest accuracy for localization of the right amygdala (p=0.02) indicating that certain brain targets may be more closely located using the MNI template as-is rather than performing individualized linear transformations.

**Figure 2.**
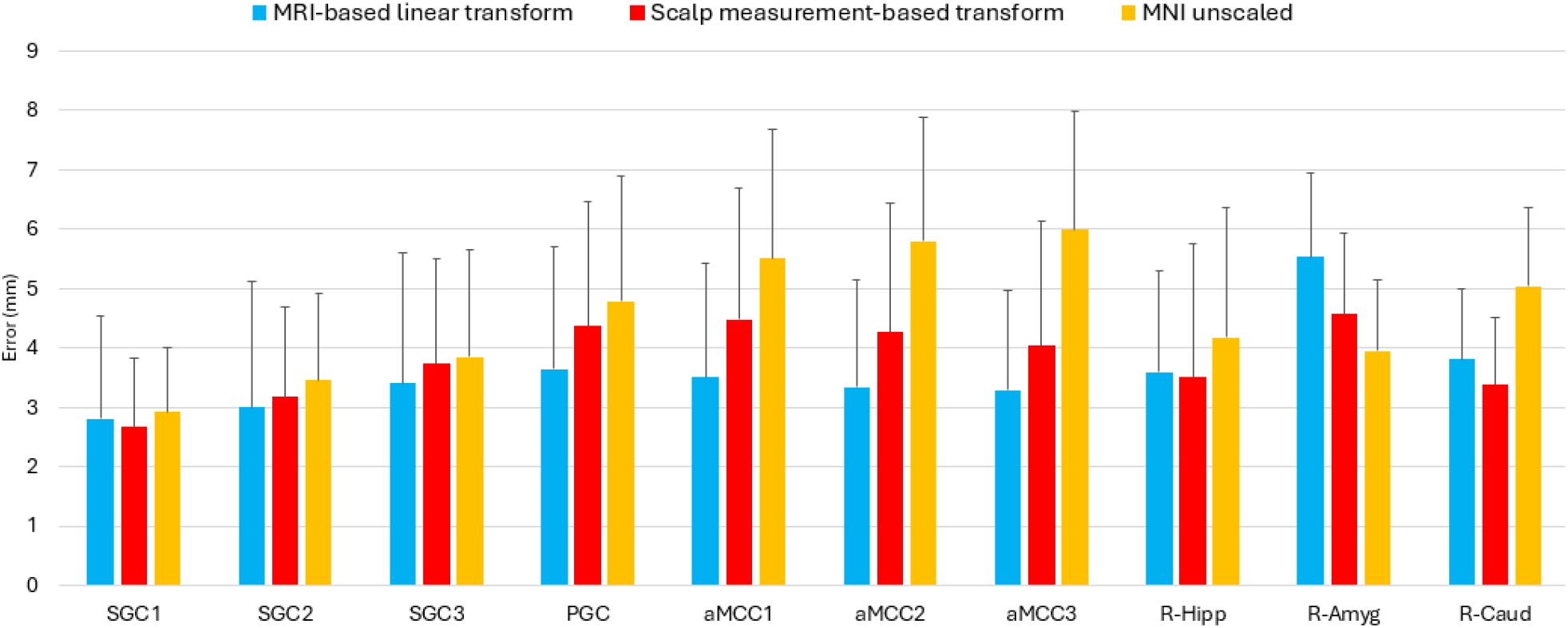
Mean errors ± SD for three different localization methods using nonlinear MRI-based transform as the ground truth.

**Figure 3.**
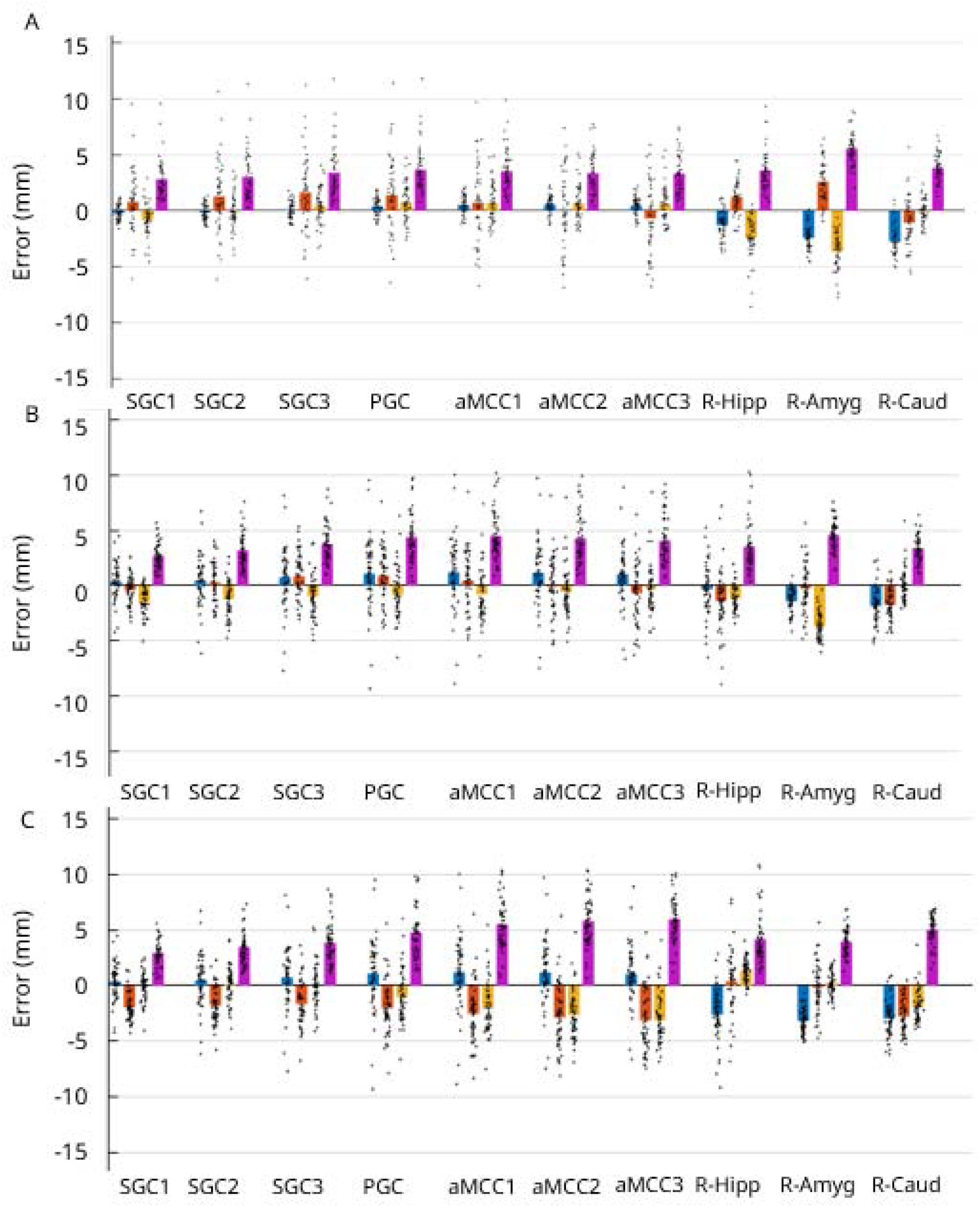
Errors in target localization for the three experimental methods in x, y, z dimensions as well as total error (R). **A:** MRI-based linear transform, **B:** Scalp measurement-based transform, **C:** MNI template unscaled. SGC: subgenual anterior cingulate cortex, PGC: pregenual midcingulate cortex, aMCC: anterior midcingulate cortex

Analysis of dimensional errors revealed that the unscaled MNI template usually overestimated the y- and z-coordinates suggesting that the MNI template is longer and taller than most brains, confirming findings by our group and others (Lee et al., 2005; Tang et al., 2010; Valter et al., 2026). For targets located on the mid-sagittal plane, the x-error for MRI-based linear localization was less than the y- and z-error. This observation is in alignment with prior reports (Riis et al., 2025) that interindividual variability in brain anatomy is greater along the y- and z-axes than along the x-axis for targets near the mid-sagittal plane.

To make this method publicly available, we provided a web-accessible tool for generating and downloading synthetic NIFTI images based on scalp measurements (tmstargets.com).

## Discussion

This study provides the first demonstration that deep brain targets can be localized using only scalp-based measurements, without reliance on neuronavigation hardware, MRI or other imaging technology. By showing that cranial landmarks alone can infer deep brain target locations, this work introduces a low-cost and accessible alternative to conventional image-guided targeting. Notably, for several targets we observed smaller localization errors than those reported for neuronavigation-based approaches using facial landmarks (Riis et al., 2025). This difference may reflect limitations inherent to facial landmarks, which are concentrated near the mid-axial and anterior regions and therefore capture only partial head geometry. In contrast, the distributed scalp measurements used here encode three-dimensional head shape. It is likely that neuronavigation-based performance would also improve with a more diverse set of anterior, posterior, and superior–inferior landmarks that better constrain 3D geometry.

Despite these advantages, the accuracy of the present method in vivo under clinical or field conditions remains untested. Practical implementation may be affected by variability in identifying anatomical landmarks and by manual scalp measurement error. Importantly, these sources of inaccuracy are not unique to MRI-free approaches; similar registration uncertainties occur in traditional neuronavigation as well (Nieminen et al., 2022). This shared limitation highlights the broader need for standardized landmark definitions, improved operator training, and consistent measurement protocols across targeting methods.

Although our approach enables accurate estimation of deep target coordinates, translating those coordinates into precise physical placement of electrodes or transducers remains a separate challenge. Without subject-specific MRI, achieving highly focal stimulation is difficult due to the complex physics of current flow and wave propagation in the head. Addressing this gap will require advances in device placement strategies and individualized modeling— directions that fall outside the scope of this initial demonstration but represent important future work.

We observed heterogeneity across brain targets. While the MRI-based linear transformation generally yielded the lowest localization errors compared to non-MRI approaches, this was not the case for all targets. These exceptions suggest that linear registration is insufficient for specific brain areas, a finding that aligns with prior research identifying a “virtual convolution” within the MNI152 template (Valter et al., 2026; Evans et al., 2012). This structural bias creates nonlinear discrepancies that cannot be corrected by linear normalization alone but require nonlinear warping to compensate for.

The framework introduced here supports multiple avenues for refinement. Potential extensions include: (1) using an unbiased or race-specific template instead of the MNI152 template (Bhalerao et al., 2018; Yang et al., 2020), (2) increasing geometric fidelity through additional or alternative cranial landmarks (Nafziger et al, 2025); (3) integrating the method with low-cost imaging or head-reconstruction technologies (Schlesinger et al., 2023, 2024; Wu, Yang, & Chang, 2022; Xiao et al., 2017) to enhance accuracy while maintaining accessibility; and (4) extending the scalp measurement-based framework to support nonlinear or more complex deformations beyond global scaling using matrix regression or machine learning methods (Nafziger et al., 2025). Collectively, these improvements may enable future MRI-free targeting systems capable not only of reliably identifying deep brain structures but also of guiding device placement with clinically meaningful precision.

